# Quantification of cell behaviours and computational modelling show that cell directional behaviours drive zebrafish pectoral fin morphogenesis

**DOI:** 10.1101/2020.08.03.235259

**Authors:** Joel Dokmegang, Hanh Nguyen, Elena Kardash, Thierry Savy, Matteo Cavaliere, Nadine Peyriéras, René Doursat

## Abstract

**Motivation:** Understanding the mechanisms by which the zebrafish pectoral fin develops is expected to produce insights on how vertebrate limbs grow from a 2D cell layer to a 3D structure. Two mechanisms have been proposed to drive limb morphogenesis in tetrapods: a growth-based morphogenesis with a higher proliferation rate at the distal tip of the limb bud than at the proximal side, and directed cell behaviors that include elongation, division and migration in a nonrandom manner. Based on quantitative experimental biological data at the level of individual cells in the whole developing organ, we test the conditions for the dynamics of pectoral fin early morphogenesis.

**Results:** We found that during the development of the zebrafish pectoral fin, cells have a preferential elongation axis that gradually aligns along the proximodistal axis (PD) of the organ. Based on these quantitative observations, we build a center-based cell model enhanced with a polarity term and cell proliferation to simulate fin growth. Our simulations resulted in 3D fins similar in shape to the observed ones, suggesting that the existence of a preferential axis of cell polarization is essential to drive fin morphogenesis in zebrafish, as observed in the development of limbs in the mouse, but distal tip-based expansion is not.

**Availability:** Upon publication, biological data will be available at http://bioemergences.eu/modelingFin, and code source at https://github.com/guijoe/MaSoFin.

**Supplementary information:** Supplementary data are included in this manuscript.

## 1 Introduction

Vertebrate limb development is a classical model system for understanding pattern formation: the process in which spatial organization of differentiated cells and tissues is generated in the embryo. Various tissue types contributing to the mature limb are derived from several embryonic tissues including the lateral plate mesoderm, the somites and the ectoderm (Christ and Brand-Saberi, 2002; Cameron and McCredie, 1982; Grim and Christ, 1993; Brand-Saberi *et al*., 1995; Huang *et al*., 2003). How the limb bud lateral plate mesoderm (LPM), which gives rise to skeletal elements and tendons, grows outward from the body trunk and acquires its particular shape remains unclear. The *in vivo* observation of the whole process and the quantification of cell behaviors underlying morphogenesis are still open challenges. The formation of the zebrafish pectoral fin bud is a model for limb development, as it is especially suited for long-term imaging owing to its external embryonic development and translucent body. The formation of the pectoral fin initiates at 18 hours post fertilization (hpf), when LPM cells condense at the prospective fin location as a flat 2D cell layer under a single layer of ectodermal cells. Over the course of the next 30 hours, the fin bud grows and forms a 3D structure. LPM cells proliferate and myoblasts coming from neighbouring somites enter the fin bud, where they will give rise to muscle bundles. At the distal tip of the fin bud, ectodermal cells align to form the so-called apical ectodermal ridge (AER) known to act as a source of molecular signaling required for the fin growth.

Over the past 40 years, the “proliferation gradient” model has been the dominant hypothesis to explain the limb bud elongation. This model suggests that a diffusible signal from the AER sets up a spatial concentration gradient. This molecule “signals the mesenchyme immediately underlying it, termed the ‘progress’ or ‘proliferative’ zone, to proliferate, resulting in directed proximodistal outgrowth” (Niswander *et al*., 1993). The AER indeed was shown to have mitogenic properties (Reiter and Solursh, 1982). On the other hand, a few studies hypothesized that directionally oriented cell behaviors drive limb elongation. Li and Muneoka (1999) demonstrated that some mesenchymal cells in the chick wing bud are capable of migrating toward an ectopic source of Fgf4 (one of the molecules produced by the AER) implanted in the center of the bud. This work directly supported the idea that mesenchymal cells could consider the Fgf gradient as a chemoattractant rather than as a mitogen.

A number of computer simulations of limb bud elongation have been produced over the last decade, most of which are in two dimensions and based on the proliferation gradient hypothesis but do not incorporate real quantitative data into their model (Dillon and Othmer, 1999; Popławski *et al*., 2007; Morishita and Iwasa, 2008). Recently, Boehm *et al*. (2010) proposed a new three-dimensional computer model based on the actual shape of the mouse limb bud captured by optical projection tomography imaging. They found that the proliferation gradient would have to be extreme from the distal end to the proximal end and subsequently too unrealistic to account for the bud shape. Similarly, Gros *et al*. (2010) concluded that uniform cell division distribution and focal regions of cell death in the chick limb bud are unlikely to be sufficient to drive the anisotropic nature of its growth. Both studies observed that cell shapes were oriented toward the nearest ectoderm, rather than distally toward the apical ectodermal ridge. They have also demonstrated the presence of filopodia suggesting active cell movement.

With improvements in *in vivo* imaging setups and data processing, a recently available complete 3D tracking of the different cell types in the early fin bud reveals their heterogeneous and complex cellular rearrangement during the transition from 2D to 3D (Nguyen *et al*., manuscript in preparation). The quantitative analysis of cell behaviors during the first 20 hours of the zebrafish pectoral fin formation are used to build a model of fin growth. The comparison between simulated and biological data highlights the pattern of cellular rearrangement underlying the pectoral fin morphogenesis.

## 2 Methods

### 2.1 Data acquisition workflow

#### 2.1.1 Zebrafish husbandry

Adult zebrafish *(Danio rerio)* were maintained at 28°C according to standard procedures as described in Kimmel *et al*. (1995). Embryos were kept at 0.3% Danieau’s medium at 28.5°C. The transgenic line *Tg(X1a.Eef11:H2B-mCherry)* was used to visualize nuclei (Recher *et al*., 2013).

#### 2.1.2 Imaging pectoral fin growth

Fin growth was monitored using the protocol described in Nguyen *et al*. (2019). Briefly, this was achieved on the upright Zeiss LSM780 confocal microscope equipped with 20x water dipping lens objective (Zeiss Objective W “Plan-Apochromat” 20x/1.0 DIC). The embryos were kept at 28.5°C and the chorion was removed using forceps prior imaging. Embryos were anaesthetized with 0.04% of tricaine methanesulfonate (Sigma) in embryo medium and mounted using agarose molds. The mounting strategy immobilized the embryo while allowing pectoral fin to develop unperturbed. The imaging plane was parallel to the plane formed by the anteroposterior (AP) and dorsoventral (DV) axes (Fig. 1a).

**Fig. 1.**
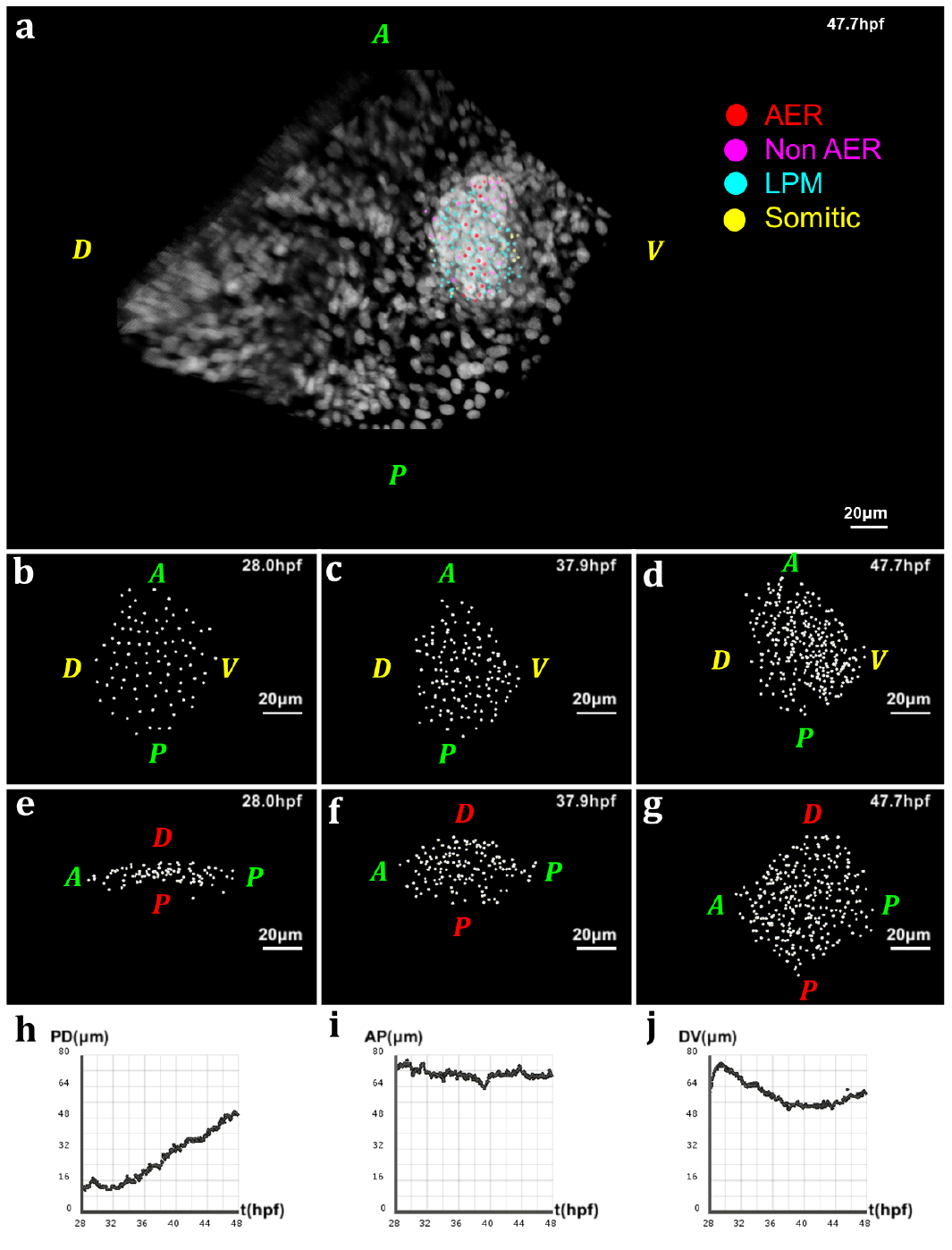
Geometry of the pectoral fin based on live imaging and image processing data. (a) 3D rendering of raw data nuclear staining at *t* = 47.7 hpf: dorsal view of the zebrafish body with detection of approximate nuclear centers of the pectoral fin cells highlighted by colored dots, where the color code depends on the cell type; scale bar: 20 μm. (b-d) After applying cell detection methods: 3D rendering of the approximate nucleus centers of LPM cells in the pectoral fin at different stages of development, respectively *t* = 28 hpf, *t* = 37.9 hpf and *t* = 47.7 hpf (AP: anteroposterior axis; DV: dorsoventral axis). (e-g) 3D rendering of the pectoral fin at the same times along the AP axis and PD (proximodistal) axis. (h-j) Evolution over time of the fin size in μm along the PD, AP and DV axes respectively. Fin expansion occurs mainly along PD. It undergoes a slight compaction along the other two axes, more pronounced along the DV axis.

#### 2.1.3 Image processing and reconstruction

The first analysis steps were performed using Fiji, an open source Javabased image processing (ImageJ) package. Raw datasets needed to undergo registration to keep the region of interest at about the same XYZ location, despite the embryos undergoing significant morphological changes during overnight imaging. To compensate for possible photobleaching, we also performed bleach correction by histogram matching. Finally, they were uploaded into the Bioemergences workflow (Faure *et al*., 2016), our online software platform integrating original mathematical methods and algorithms to perform image filtering, nucleus center detection, and cell tracking (Fig. 1b-g). The outcome of cell detection was then manually validated using Mov-IT, an interactive visualization and editing tool complementing Bioemergences.

### 2.2 Tracking zebrafish pectoral fin growth along PD, AP and DV axes

The quantitative analysis required for this work necessitated that we track the fin’s dynamics along its main axes: proximal-distal (PD), anterior-posterior (AP), dorsal-ventral (DV). Precise tracking of the fin’s size along is a challenge due to the fact that the embryo moves during imaging. Embryo movements mean that its main axes do not always maintain a static orientation. This in turn poses the problem of precise identification of the fin’s main axis for every time point. To tackle this issue, at each time point, we computed new orientations for the PD, AP, DV axes as functions of both the cloud of points (cell centers) at the current time and their orientations at the previous time step. First, we applied the Principal Component Analysis (PCA) algorithm to the cell centers in order to determine the three main directions of the point cloud. Then, we compared each of PCA output axes with the PD axis at the previous time step and kept the most parallel one as our new PD axis. Given that PCA does not keep track of the orientation at previous time steps, the basis formed by the two remaining PCA axes could be significantly rotated compared to the previous AP-DV basis. To correct this issue, we projected onto the plane formed by these two vectors the AP and DV axes computed at the previous time steps, and considered these projections to be our new AP and DV axes. When the movement of the fin as a whole between two time steps is not significant enough to change the orientation of the axes, we keep the orientations computed in the previous time step. Here, noticing that the fin as whole, although still growing, is relatively stable as from about 43.3hpf, we stopped computing new axes orientations as from about that time point.

### 2.3 Computational model

In order to test data-derived hypothesis, we turned to computational modelling. When it comes to spatially explicit simulations of biological development, several computational approaches stand out. Depending on the biological realism they include, the spatiotemporal scale they capture, or the nature of variables they manipulate, models exhibit different properties and suit different purposes (Van Liedekerke *et al*., 2015; Marin-Riera *et al*., 2015). Here we set our choice on the family of center-based models (CBM) due to their simplicity and compatibility with our data. In CBM, cells are described by simple geometrical shapes whose representation can be reduced to their centers. These models assume that cell trajectories in space can be assimilated to the motion of particles, which are governed by an equation of motion. CBMs have been used extensively to study the development of multiple organisms (Delile *et al*., 2017; Villoutreix *et al*., 2016; Germann *et al*., 2019; Van Liedekerke *et al*., 2018).

We used a simplified mechanistic formulation of a CBM approach to simulate pectoral fin growth (Fig. 2a). Cells are subjected in their center to forces governing their behavior. Similar to Delile *et al*. (2017), we distinguished two main types of forces acting on a cell.

**Fig. 2.**
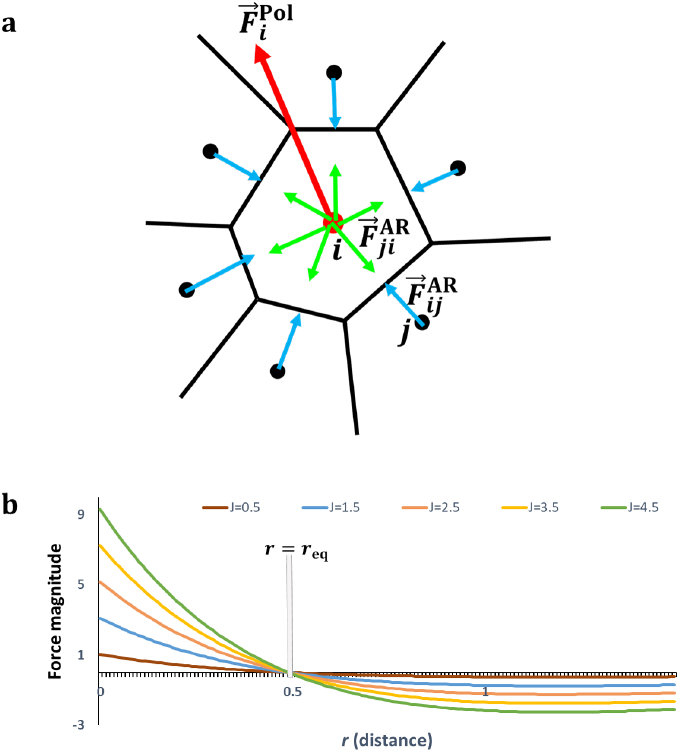
Center-based computational model of multicellular dynamics. (a) Schema of a local cell neighborhood and the abstract forces on cell centers. 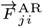 is the passive attractionrepulsion force exerted on a cell *i* by a cell *j*. 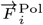 is the active migration force driven by the cell’s polarity (specified in Section 3.4). (b) Plot of the Morse force profile (derivative of the Morse potential) defining 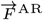, for different parameter values. This curve presents two regimes: a positive regime (attraction) below an equilibrium distance *r*_eq_ and a negative regime (repulsion) above.

On the one hand, passive attraction-repulsion (AR) forces 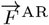 regulate interactions between cells and their local neighborhoods. In the literature, these forces often derive from elastic potential mimicking linear or non-linear springs (Drasdo and Loeffler, 2001; Basan *et al*., 2011). Here, we followed this rule and derived AR forces from a “Morse potential”, a curve exhibiting a quadratic minimum framed by vertical and horizontal asymptotes (see its derivative in Fig. 2b). Moreover, we also defined a neighborhood for each cell by computing the 3D Delaunay tetrahedralization of the system. Two cells are deemed neighbors if their centers belong to the same tetrahedron.

On the other hand, cells also exhibit an active migration force which shapes their motion. Informed by empirical evidence, we constructed a polarity-driven migration force 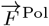 governing cells’ intrinsic mechanics (explained in Results, Section 3.4). Finally, we neglected the effects of inertia due to a low Reynolds number (Odell *et al*., 1981; Delile *et al*., 2017), and only considered viscosity-driven friction via a constant coefficient λ. Altogether, the equation of motion for a cell *i* with neighborhood 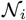 reads:

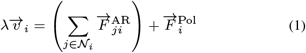

where 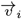 is the velocity of cell *i*.

In CBM, cell divisions are often simulated by adding daughter cells to the system at appropriate positions, notably in the neighborhood of the mother cell. Here, we assumed that cells divide along their long axis and added new cells such that mother and daughter cells were aligned along this axis. Furthermore, we dealt with cell cycles by setting a global cycle period for all cells, and assigning random initial phases to individual cells. Using this model, we simulated zebrafish pectoral fin morphogenesis starting from the initial cell arrangement provided by the imaging data.

## 3 Results

### 3.1 Zebrafish pectoral fin morphogenesis is proximal distal oriented

Using the approach described in the methods, we computed the pectoral fin’s main axes for each time point. Then, we proceeded with calculating the size of the fin along each direction (Fig. 1h-j). Our data suggests that the fin expands principally along the PD axis with a quasi-linear slope, after an initial oscillatory behavior (Fig. 1h). This result aligns with previous observations which has consistently shown that a common property of limb development in vertebrates is the distal orientation of their growth (Hopyan, 2017; Boehm *et al*., 2010). Furthermore, in this dataset, while fin’s length along the AP axis oscillates somewhat, no significant overall change is recorded (Fig. 1i). Along the DV axis, however, the fin seems to contract slightly over the length of development (Fig. 1j), but seems to recover in the latest time steps.

### 3.2 Distal tip-based growth does not account for zebrafish pectoral fin morphogenesis

We sought to determine the role of proliferation in zebrafish pectoral fin growth. For this, we computed for every time point of development the bounding box encapsulating the pectoral fin. Then, we discretized this bounding box using the same volume unit everywhere. Next, we calculated the cumulative number of cell divisions in each volume unit of that space over the duration of fin growth. In order to understand the distribution of proliferation in 3D space, we proceeded with plotting the marginal distributions of cumulative divisions along the three main axes of the pectoral fin (Fig. 3). To ensure that our observations were not mere features of a single developing fin, we applied the same analysis to a supplementary dataset, consisting, as the main one, to live imaging of a developing zebrafish pectoral fin (Fig. S1).

**Fig. 3.**
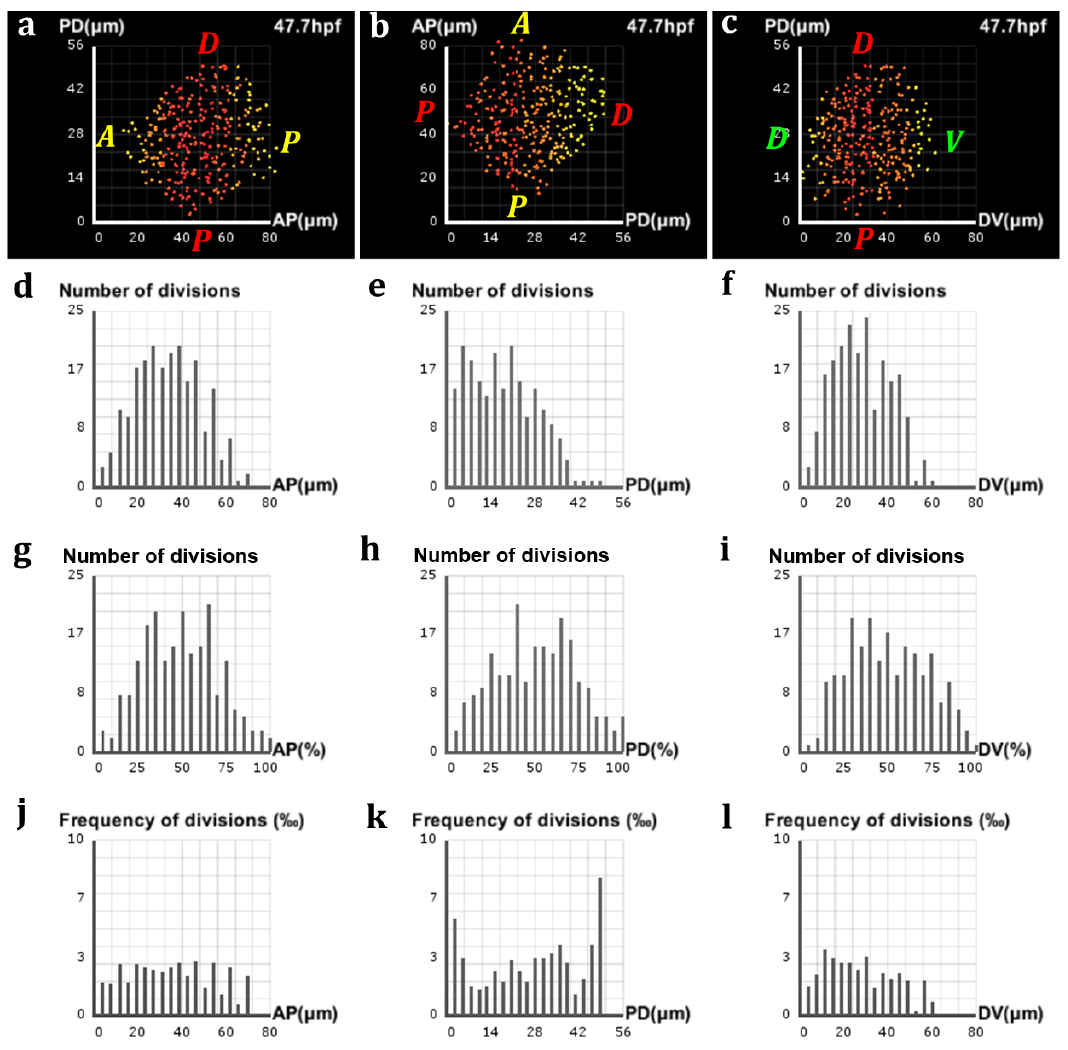
Analysis of proliferation in the zebrafish pectoral fin. (a-c) Frequencies of divisions along the AP, PD and DV axes respectively, highlighted by a yellow-red color gradient coding for differences in proliferation rates across the fin. (a,c) The preponderance of red at the center of the fin shows where the bulk of cell divisions takes place, with only a few of them occurring near the lateral surfaces (yellow). (b) A decreasing gradient of proliferation rates from the proximal pole to the distal tip characterizes the PD axis. (d-f) Marginal distributions of proliferation along the AP, PD and DV axes respectively, expressed in numbers of cells with respect to the absolute distance in μm along the axis. (g-i) Same distributions with respect to the relative distance on the axis. (j-l) Same distributions expressed in proportions of cells with respect to the absolute distance.

Marginal distributions of proliferation along the AP and DV axes show that during pectoral fin morphogenesis, the bulk of proliferation is concentrated at the center of the fin, while only a few divisions are observed near the lateral surfaces (Fig. 3a,c, Fig. S1a,c). Differential behaviors of cells based on their location is a well-established biological mechanism, reminiscent of the so-called “French flag” model of Wolpert’s positional information (Wolpert *et al*., 2015). Furthermore, histogram plots of these marginal distributions may suggest that proliferation along the AP and DV axes can be assimilated to Gaussian processes (Fig. 3d-f, Fig. S1d-f). Although it could be expected that such behavior facilitates the development of the fin toward its known shape, it is not clear whether it is sufficient to drive this growth. However, it is also likely that the accumulation of cell division in the inner volume of the fin (Fig. 3g-i, Fig. S1g-i) is merely a consequence of the fin’s geometry, namely its overall conic shape, favoring higher number of cells in the middle than near the lateral surfaces. This interpretation seems to be favoured by the data in Fig. 3j,l (also Fig. S1j,l), which present quasi-uniform histograms of number of divisions per time step over the length of AP and DV axes.

Along the PD axis, the marginal distribution of proliferation shows a decreasing gradient of cell division from the proximal pole to the distal tip of the fin (Fig. 3b, Fig. S1b). This simply translates into the fact that proximal layers of the fin, which form early, host more divisions than distal regions, which develop later. This observation stands in contradiction with the growth-based morphogenesis hypothesis that stipulates higher proliferation rates at the distal tip of the fin. Hence, this result suggests that growth-based morphogenesis might not be the main drive for zebrafish fin pectoral morphogenesis.

### 3.3 Zebrafish pectoral fin cells exhibit preferential directional behaviors

Next, we looked whether cells exhibited peculiar behaviors along preferential directions that could influence the shaping of the zebrafish fin. To this goal, we decided to analyse the dynamics over time of the elongation axis of each cell. We determined the elongation axis of a cell *i* by computing the direction of maximum variance of the cloud of points consisting of cell *i* and its Delaunay neighborhood 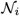. This direction was given by the eigenvector corresponding to the maximum eigenvalue of the covariance matrix of 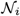. We denote this vector by 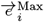, which we also consider to be the polarity vector of the cell (Fig. 4a).

**Fig. 4.**
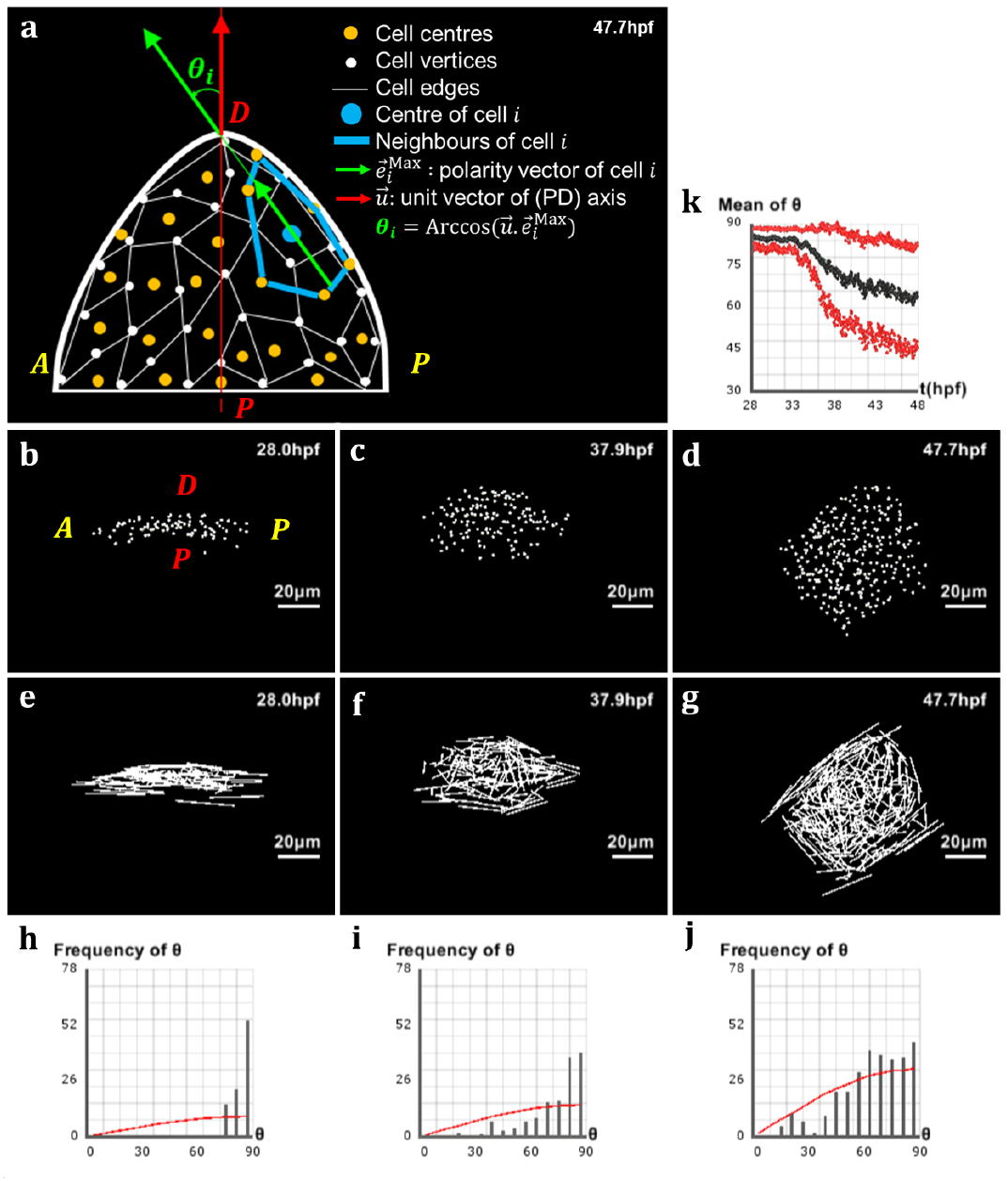
Analysis of directional cell behaviors in the zebrafish pectoral fin. (a) Schematics in 2D of the method used to analyse directional cell behaviors: foreach cell *i, θ_i_* denotes the polarity angle that this cell forms between its elongation axis 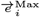 (extracted from the maximum eigenvalue of the covariance matrix of its neighborhood 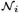) and the PD axis 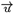. (b-d) Lateral view of the pectoral fin at different stages of development, respectively *t* = 28.0 hpf, *t* = 37.9 hpf and *t* = 47.7 hpf. (e-g) Vector field of the cells’ elongation axes 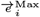 in the pectoral fin at the same stages. (h-j) Distribution of the polarity angles *θ_i_* of the cells in the pectoral fin at the same stages, compared with the standard distribution of random angles formed by two arbitrary vectors in 3D (red curve). (k) Evolution over time of the average polarity angle 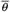 of the fin cells ± its standard deviation Δ*θ* shown in red.

Having computed the elongation axis of each cell through every time point of development, we observed that cells at initial stages are elongated perpendicularly to the PD axis. We further noticed that, during development, cells gradually bring their long axis in closer alignment to the PD axis (Fig. 4e-g). In order to confirm this qualitative observation, we measured the angle that cells form via their long axis with the PD axis of the pectoral fin, and called it the “polarity angle” with notation *θ_i_* (Fig. 4a). At the initial time point, the distribution of polarity angles was clustered around 90°, confirming the previous observation that cells were elongated perpendicular to the PD axis (Fig. 4h). During development, this distribution spread in a nonrandom way between 0 and 90°, where the average polarity angle decreased toward a value of 60°, meaning that cells exhibited preferential directionality by orienting their long axis toward the PD axis (Fig. 4i,j). To ensure that our observations were not mere features of a single developing fin, we applied the same analysis to a different dataset which lead to a similar dynamics (Fig. S2).

### 3.4 Directional cell behaviors are essential to drive zebrafish pectoral fin morphogenesis

We wanted to find out whether directional behaviors of cells were sufficient to drive fin morphogenesis. In the previous analysis, we observed that cells tended to align their long axis in the direction of the PD axis, eventually forming an average polarity angle 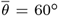. We noticed that such behavior could explain the overall conic shape of the fin. Based on this observation, we designed the “polarity force” term 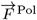 of the 3D model as follows. First, we defined a global “polarity energy” of the cell population, denoted by *H*^Pol^. Models in which forces derive from problem-specific global energy have been used in different contexts such as cell sorting, molecular signaling, or epithelial morphogenesis in the developing *Drosophila* (Fletcher *et al*., 2014; Osborne *et al*., 2017). Germann *et al*. (2019) also use an energy term conjointly with CBM by defining a tissue polarity potential and an apicobasal polarity to distinguish between epithelial and mesenchymal tissues.

Here *H*^Po1^ is expressed over all *N* cells by:

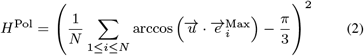

where 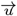 denotes the unit vector of the PD axis. Although *H*^Po1^ is defined globally, individual cells contribute to this potential only to the extent of their local neighborhood. This energy was designed such that its minimum corresponds to 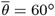. Then, we set the polarity force to be proportional to the opposite of the gradient of *H*^Pol^ with respect to each cell:

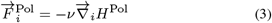

Having defined these laws governing cell interactions, we proceeded to simulating limb morphogenesis. In order to highlight the influence of the polarity force, we used a uniform cell cycle period with a random initial phase for each cell. Our virtual limb featured similar properties to the imaged limb (Fig. 5). Driven by the polarity force 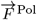 and constrained by the elastic force 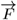^AR^ defined in Section 2.3, cells moved and reshaped their neighborhoods to minimize the polarity energy 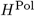, resulting in 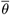 effectively decreasing throughout development toward 60° (Fig. 5n), in a way similar to the real fin. The polarity angle distribution *θ_i_*, which clustered around 90° at the initial time point, progressively spread between 0 and 90° (Fig. 5h-j). Another result is that this force restricted the fin growth toward the distal pole, as observed in development. Furthermore, over the same period of time as in our dataset, the virtual fin grew to a size comparable to that of the real fin, from about 12.71 μm to about 52.94 μm (Fig. 5k), presenting over imaged fin an increase of just below 7%. Finally, our simulated fin acquired a global conic shape similar to that of the real fin (Fig. 5b-d,e-g). Taken together, these results suggest that directional cell behaviors, in particular alignment toward the PD axis of the zebrafish pectoral fin, could be an essential drive for its morphogenesis.

**Fig. 5.**
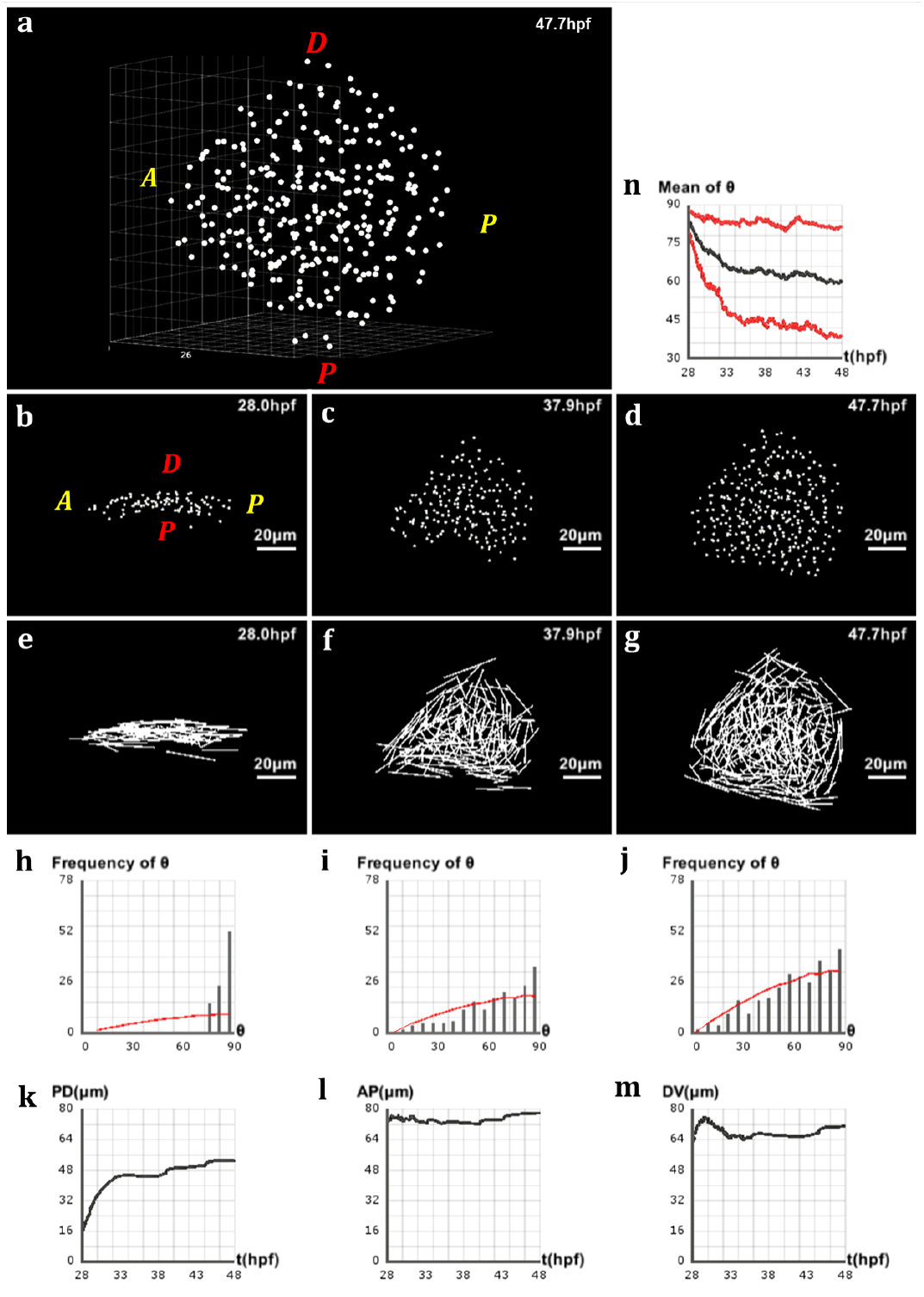
Simulation of pectoral fin morphogenesis based on directional cell behaviors. Values of the equation parameters: λ = 0.2, *ν* = 1. (a)3D view of the simulated fin at the final stage *t* = 47.8 hpf. (b-d) Lateral view of the simulated fin at different stages of development, respectively *t* = 28.0 hpf, *t* = 37.9 hpf and *t* = 47.7 hpf. (e-g) Vector field of the cells’ elongation axes 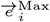 in the simulated fin at the same stages. (h-j) Distribution of the polarity angles *θ_i_* of the cells in the simulated fin at the same stages, compared with the standard distribution of random angles formed by two arbitrary vectors in 3D (red curve). (k-m) Evolution over time ofthe simulated fin size in μm along the PD, AP and DV axes respectively. We observe roughly the same behavior as the real fin in Fig. 1h-j. (n) Evolution over time of the average polarity angle 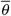 of the simulated fin cells ± its standard deviation Δ*θ* shown in red. This curve is more scattered than Fig. 4k.

## 4 Discussion

The question of how vertebrates make their limbs is a fascinating problem in embryology that has been widely investigated across multiple species (Tickle, 2015). Although the study of molecular patterns underlying this morphogenesis has provided rich insights, the cellular basis of limb formation has not been completely elucidated. The availability of *in toto* imaging with resolution at the single cell level provides quantitative data for cells’ behavior along their trajectory. New methods need to be developed to analyze such data, to use it to feed realistic models and evaluate hypotheses through a quantitative comparison between *in vivo* and *in silico* data. This work brings insights into the development of the zebrafish pectoral fin using quantitative analysis of imaging data and computational modelling.

Here, we investigated zebrafish pectoral fin development under the prism of the two dominant hypotheses of cellular behavior during limb growth. On the one hand, we analyzed proliferation behaviors in different regions of the fin, and found that proliferation gradients could not account for the observed growth. On the other hand, analysis of cellular elongation directions showed that cells tended on average to lower the angle they formed with the PD axis via their long axis, an indication of preferential polarity. To test this hypothesis, we formulated a simple mechanistic model of cells and simulated pectoral fin morphogenesis.

Our model of fin development based on quantitative biological data derived from live imaging and image processing accounts for the directional growth and the shape of the fin. However, due to some differences in comparison with imaging data, it also helps refine hypotheses concerning additional constraints that were not integrated here. The simulated fin does not exhibit the same amplitude of slight compaction along the DV axis as the one measured in the biological data. In addition, the simulated fin shows a first phase of fast growth not quite observed in the zebrafish where the fin growth rate is closer to linear. We hypothesize that constraints imposed by the outside cell layers (i.e. ectodermal layer and enveloping layer) also contribute to the regulation of the fin growth and shaping, as observed in chick limb morphogenesis (Popławski *et al*., 2007). Constraints from the outside cell layers, or other sources including for example electrical fields (Hopyan, 2017), could also play a role in the emergence of the cell polarization axes implemented in our model. We considered the contribution of somitic cells that invade the fin bud during the time course of our observation as neutral regarding the overall growth and shaping. Both an *in vivo* and *in silico* experimentation are needed to support this view.

The biomechanical aspects privileged in our study are part of a complex interplay of genetic and mechanical cues characteristic of biological development. If the response of LPM cells during early fin development may mainly involve mechanical transmission, it is certainly required to integrate genetic and molecular interactions as well in the transformation of the ectoderm that leads to shape the AER. Further studies may be integrate these mechanisms for more insights into zebrafish pectoral fin morphogenesis.

## Acknowledgements

We would like to thank Lianxiu Han, Moi Hoon Yap, Monique Frain and Antoine Gaget for useful conversations related to this work.

## Supplementary figures

**Fig. S1.**
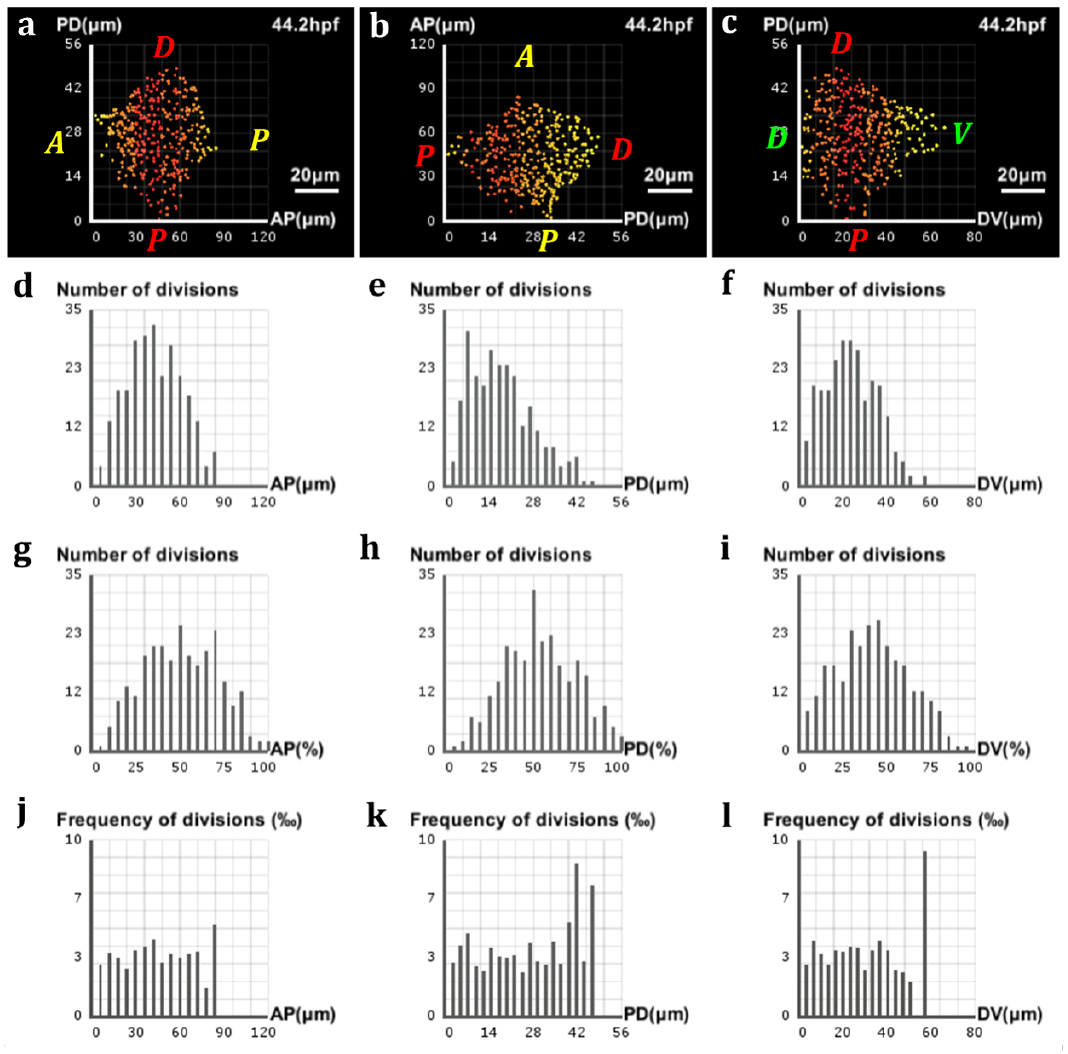
Analysis of proliferation in the zebrafish pectoral fin (supplementary dataset). (a-c) Frequencies of divisions along the AP, PD and DV axes respectively, highlighted by a yellow-red color gradient coding for differences in proliferation rates across the fin. (a,c) The preponderance of red at the center of the fin shows where the bulk of cell divisions takes place, with only a few of them occurring near the lateral surfaces (yellow). (b) A decreasing gradient of proliferation rates from the proximal pole to the distal tip characterizes the PD axis. (d-f) Marginal distributions of proliferation along the AP, PD and DV axes respectively, expressed in numbers of cells with respect to the absolute distance in μm along the axis. (g-i) Same distributions with respect to the relative distance on the axis. (j-l) Same distributions expressed in proportions of cells with respect to the absolute distance.

**Fig. S2.**
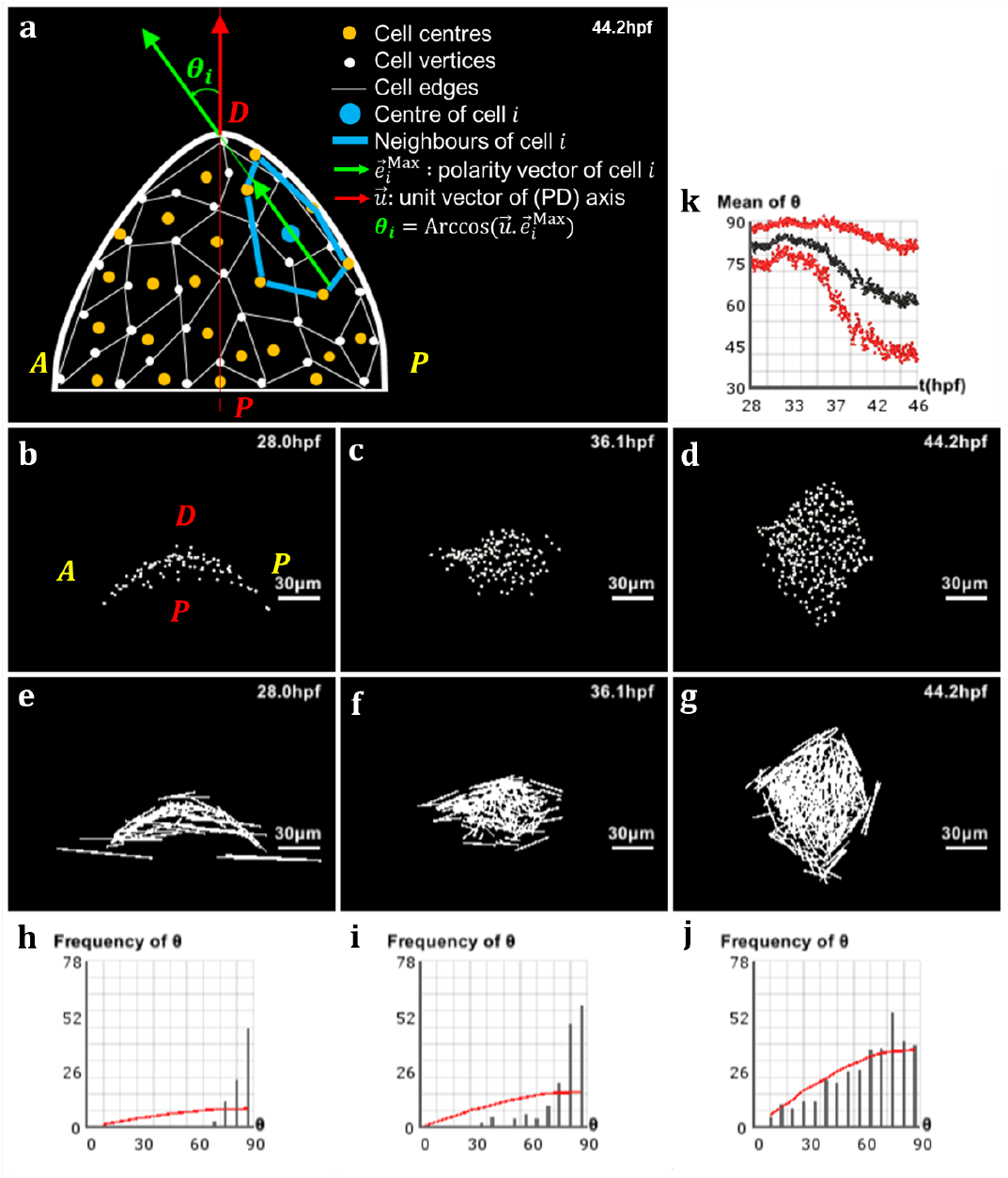
Analysis of directional cell behaviors in the zebrafish pectoral fin (supplementary dataset). (a) Schematics in 2D of the method used to analyse directional cell behaviors: for each cell *i, θ_i_* denotes the polarity angle that this cell forms between its elongation axis 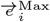 (extracted from the maximum eigenvalue of the covariance matrix of its neighborhood 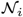) and the PD axis 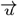. (b-d) Lateral view of the pectoral fin at different stages of development, respectively *t* = 28.0 hpf, *t* = 36.1 hpf and *t* = 44.2 hpf. (e-g) Vector field of the cells’ elongation axes 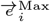 in the pectoral fin at the same stages. (h-j) Distribution of the polarity angles *θ_i_* of the cells in the pectoral fin at the same stages, compared with the standard distribution of random angles formed by two arbitrary vectors in 3D (red curve). (k) Evolution over time of the average polarity angle 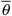 of the fin cells ± its standard deviation Δ*θ* shown in red.

